# Deep Learning-Assisted Evaluation of Laryngeal Mobility in a Rat Model

**DOI:** 10.64898/2026.02.26.708292

**Authors:** Amin Mirzaaghasi, Eric M. Smith, Ashley E. Kita

## Abstract

Vocal fold mobility is essential for normal laryngeal function and is often compromised following recurrent laryngeal nerve (RLN) or superior laryngeal nerve injuries. In a rat model of unilateral (RLN) injury, we quantitatively evaluate laryngeal mobility using advanced computer vision techniques. Adult male Long-Evans rats underwent direct laryngoscopy before and after RLN injury at the level of the 5th tracheal ring. High-resolution video recordings were analyzed with the open-source deep learning framework Social LEAP Estimates Animal Poses (SLEAP) to track key laryngeal landmarks. The displacement of the left and right arytenoid processes relative to the anatomical midpoint was measured frame-by-frame. Mean differences and 95% CI were computed for each arytenoid. A mean difference threshold of 0.42 differentiated laryngeal asymmetry from symmetry. This method offers a quantitative method of assessing laryngeal symmetry in a rat model.

## Introduction /Background

Vocal fold mobility is critical for laryngeal functions such as phonation, swallowing, and respiration, and conditions affecting this motion can significantly impact quality of life [1,2]. Animal models, particularly rats, are essential in preclinical investigations of laryngeal pathologies [3,4]. A rat model of RLN crush injury is widely used to study nerve injury and evaluate interventions for functional recovery [4,5]. However, recovery monitoring has traditionally relied on euthanizing cohorts of rats at multiple time points to estimate recovery timing [6]. While informative, this approach is imprecise and fails to account for individual variation.

Currently, laryngoscopy is the gold standard for assessing vocal fold function in humans and animals [7,8]. In rats, however, it is challenging due to their small size and anesthetic difficulties [9,10]. Ketamine/xylazine protocols have been used to avoid gas anesthetics, but they carry high mortality and often prolong anesthesia beyond what is needed for laryngeal visualization [11]. These factors limit repeat assessments in longitudinal studies. Alternatives such as laryngeal ultrasonography have shown promise, with Dewan et al. reporting 100% sensitivity and specificity in detecting cord mobility after crush injury [10–12]. Digital otoscopes have also been adapted for rat laryngoscopy and intubation to address size limitations of traditional devices [13]. Here, we describe a method to detect, track, and quantify subtle changes in vocal fold mobility, enabling monitoring of recovery and evaluation of new treatments for laryngeal disorders, thereby addressing a critical need in this field [3,4].

### Animal Model and Surgical Procedure

Adult Long Evans rats (n = 4, ∼280 g, equal gender) were anesthetized with isoflurane induction and ketamine-xylazine injection. Baseline laryngeal movement was recorded using a digital otoscope with iPhone integration (Ear Wax Removal Tool, Amazon, Seattle, WA). Physiological parameters were monitored using rectal temperature probe, heating pad, and pulse oximetry. The anterior neck was prepped, and a vertical midline incision was made to expose the trachea. The right recurrent laryngeal nerve (RLN) was identified and crushed at the level of the 5th tracheal ring for 60 seconds using Jeweler’s forceps (Dumont biology, Switzerland) (Fig. 1). Post-injury laryngoscopy confirmed grossly diminished right arytenoid mobility in all animals, and post-injury videos were recorded. Rats were euthanized after the procedure. All experiments were approved by the UCLA Animal Research Committee (ARC 2021-005).

**Figure 1.**
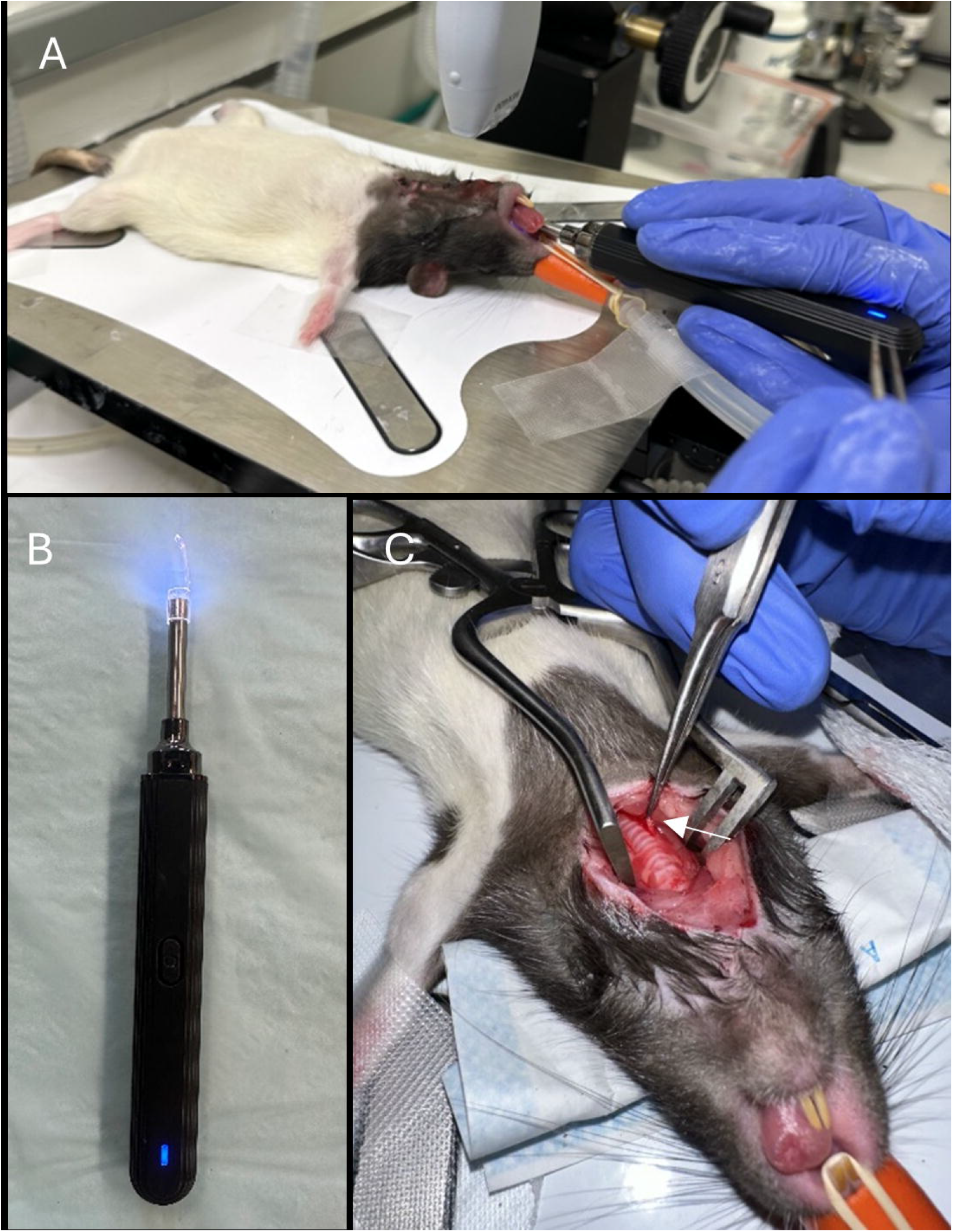
Positioning and equipment for video laryngoscopy and RLN crush surgery (A) Video laryngoscopy position (B) Human otoscope used for laryngoscopy (C) RLN crush injury, arrow shows 5th tracheal ring.

### Video Recording and Analysis

Two video segments of approximately one minute were recorded per animal pre- and post-crush injury. A total of 16 video recordings were obtained for analysis. Videos were then manually trimmed to include segments with stable laryngeal visualization.

To quantify laryngeal mobility, we developed a computer vision module using Social LEAP Estimates Animal Poses (SLEAP) [13]. Trimmed videos were imported and 20 randomly selected frames per video were manually labeled at predefined laryngeal landmarks for training. After 35,000 iterations, the model produced fully labeled videos and coordinate data (.csv files). Displacements of the left (LV2) and right (RV2) arytenoid cartilages were calculated as the distance from each arytenoid to an anatomical midpoint between the left (LC) and right (RC) corners of the larynx (Fig. 2). These frame-by-frame displacements were used to quantify arytenoid mobility. We hypothesized that RLN crush injury would result in reduced arytenoid movement on the ipsilateral side as compared to the uncrushed side, as quantified by this laryngoscopy-based tracking method [1,4].

**Figure 2.**
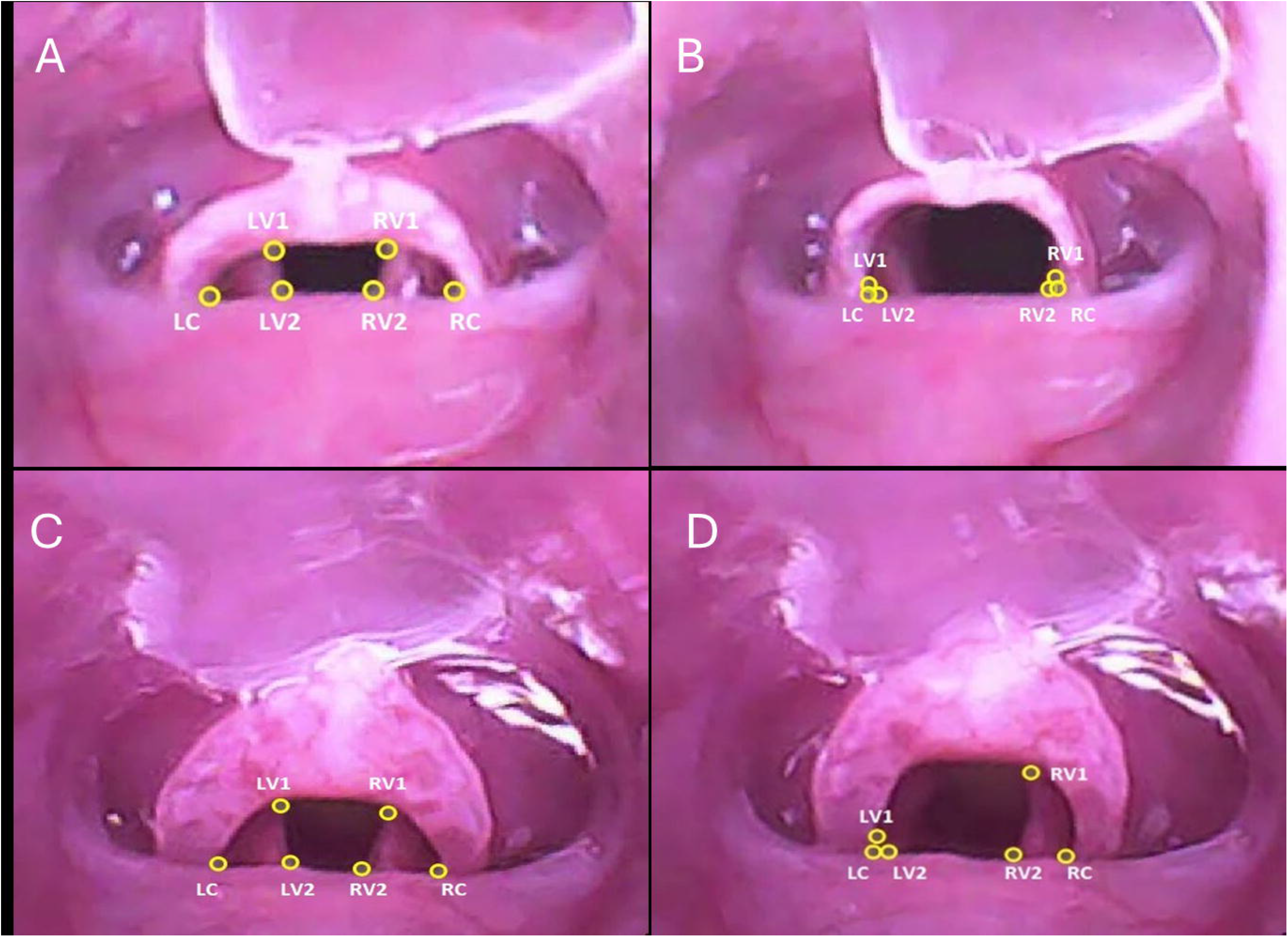
Rat larynx showing arytenoids pre-crush in adducted (A) and abducted (B) positions, and post-crush in adducted (C) and abducted (D). The right arytenoid’s paramedian position indicates successful unilateral RLN crush injury.

### Development of a Threshold to Determine Laryngeal Mobility in Rats

After determining the number of cycles of arytenoid opening and closing in each video based on graphs of X and Y coordinate displacement of RV2 and LV2 over time, the displacement values for each cycle were determined for the left and right arytenoid.

After normalizing all displacement values for an individual video to the average displacement of the uncrushed arytenoid (left) for each video to allow comparison between videos, mean differences between right and left arytenoid displacements were then calculated. Mean differences and associated standard errors were used to construct 95% CI. Analyses were performed using Microsoft Excel v2408. Continuous variables are presented as mean ± standard error. Group comparisons were conducted using standard parametric tests (Student’s *t* test). A significance level of *p* < 0.05 was adopted for all comparisons. Confidence intervals (CI) for mean differences were calculated using a t-distribution.

The difference between left and right arytenoid displacement in pre-crush videos was expected to be non-significant. This was true for most, but not all videos. The difference between left and right arytenoid displacement in post-crush videos was expected to be significant, and this was true for all but one video (Supplemental Table 1). After normalizing each video’s displacements to the average uncrushed (left) arytenoid, right–left mean differences and their standard errors were calculated and used to derive 95% confidence intervals via the t-distribution (mean ± t_{df,0.975}·SE, with SE = SD/√n) for each video; the overall mean of the differences between the pre- and post-crush videos (0.42) was taken as the reference, with values <0.42 indicating symmetric laryngeal movement and values >0.42 indicating unilateral reduced movement (Fig. 3).

**Figure 3.**
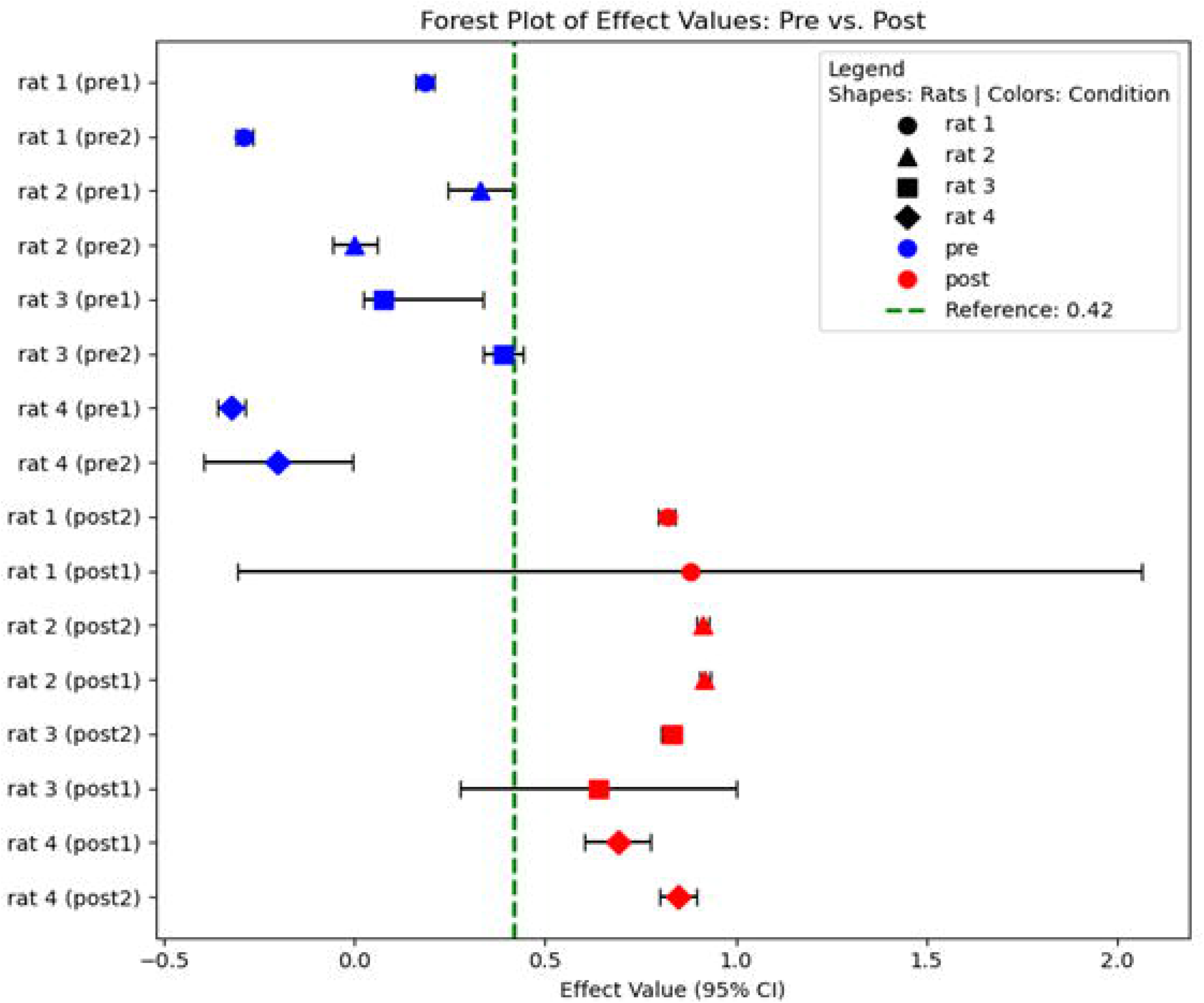
Forest plot of displacement values (95% CI) pre-crush (blue) vs. post-crush (red). Markers show individual rats; green dashed line marks mean reference distinguishing laryngeal symmetry from asymmetry.

## Conclusion

This quantitative approach allows identification of unilateral laryngeal weakness in a rat RLN injury model, using a mean displacement difference threshold of 0.42 between arytenoids to distinguish impaired from symmetric mobility. This approach describesa useful tool for future studies tracking laryngeal mobility in response to injury and/or treatment.

---

Supplemental Table 1. Individual video t test values.

