## Supplementary figures and images for "Deep Learning-Assisted Evaluation of Laryngeal Mobility in a Rat Model"

### Supplemental Table.1

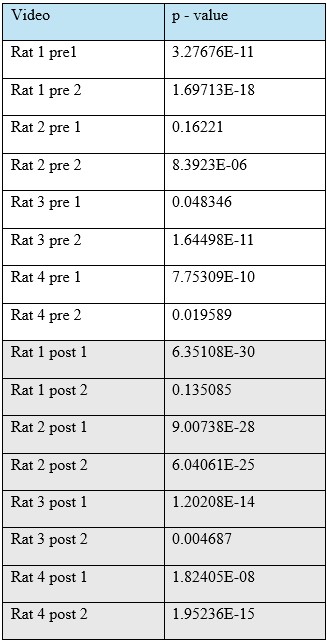
